# Spatiotemporal shifts in the role of floral traits in shaping tropical plant-pollinator interactions

**DOI:** 10.1101/2020.10.16.342386

**Authors:** Yannick Klomberg, Robert Tropek, Jan E.J. Mertens, Ishmeal N. Kobe, Jiří Hodeček, Jan Raška, Nestoral T. Fominka, Daniel Souto-Vilarós, Štěpán Janeček

**Affiliations:** Department of Ecology, Faculty of Science, Charles University, Viničná 7, 12844 Prague, Czechia; Naturalis Biodiversity Center, Darwinweg 2, 2233CR Leiden, The Netherlands; Institute of Entomology, Biology Centre, Czech Academy of Sciences, Branišovská 31, 37005 České Budějovice, Czechia; Department of Zoology and Animal Physiology, Faculty of Science, University of Buea, Buea, Cameroon

**Keywords:** Foraging behaviour, Afrotropics, Plant-Pollinator Interactions, Mount Cameroon National Park, Pollination syndrome, pollinator predictability

## Abstract

The pollination syndrome hypothesis predicts that plants pollinated by the same pollinator group bear convergent combinations of specific floral traits. Nevertheless, some studies have shown relatively low predictive power for these floral trait combinations. This discrepancy may be caused by changes in the importance of specific floral traits for shaping interactions under different environmental conditions and for different pollinator groups. To test this, we studied pollination systems and floral traits along an elevational gradient on Mount Cameroon during wet and dry seasons. Using Random Forest models, allowing the ranking of traits by significance, we demonstrated that some floral traits are more important than others in shaping interactions and that these traits predict pollinators relatively well. However, the distribution and importance of traits varies under different environmental conditions. Our results imply the need to improve our trait-based understanding of plant-pollinator interactions to better inform the debate surrounding pollination syndrome hypothesis.

## Introduction

The importance of floral traits for plant-pollinator interactions has been apparent since the 18^th^ century (Knuth, 1906; Müller, 1883; Sprengel, 1793). Darwin placed the origin of floral traits into the modern evolutionary framework (Darwin, 1859, 1862) and during the 19th and 20th century other scholars followed by suggesting various floral trait classifications according to their adaptive relationships to particular pollinator groups. These efforts resulted in an influential ecological and evolutionary hypothesis, the pollination syndrome hypothesis (Delpino, 1874; Faegri & van der Pijl, 1979; Fenster, Armbruster, Wilson, Dudash, & Thomson, 2004; Waser, 2006). It is defined as a set of convergent floral traits (e.g. colour, shape, odour or production and display of floral rewards) evolved to attract a particular group of pollinators (Faegri & van der Pijl, 1979).

Despite such a long research history, the pollination syndrome hypothesis has been questioned in recent decades. One of the main reasons is that community wide studies exploring complex plant-pollinator networks demonstrated a higher level of generalisation in pollination systems and lower predictability of pollinators based on floral traits than previously expected (S. D. Johnson & Steiner, 2000; Ollerton et al., 2009; Waser, Chittka, Price, Williams, & Ollerton, 1996). Recent empirical efforts have provided evidence both supporting (e.g. Fenster, Reynolds, Williams, Makowsky, & Dudash, 2015; Hargreaves, Johnson, & Nol, 2004; Rosas-Guerrero et al., 2014; Vandelook et al., 2019) and contradicting (e.g. Blüthgen, Menzel, Hovestadt, Fiala, & Blüthgen, 2007; Ollerton et al., 2009; Paudel, Kessler, Shrestha, Zhao, & Li, 2019; Rocha, Domingos-Melo, Zappi, & Machado, 2020; Wang, Wen, Qian, Pei, & Zhang, 2020) the validity of the pollination syndrome hypothesis. Researchers also demonstrated that pollination systems can show parallel adaptations to multiple pollinator groups (Dellinger, Scheer, et al., 2019), and thus suggested reclassification of some pollination syndromes (Dellinger, Chartier, et al., 2019), or proposed that pollination syndrome theory can be improved by other concepts, like optimal foraging theory or evolution stable strategy (Pyke, 2016).

At the same time detailed studies on individual traits “packed” within pollination syndromes have shown that we do not fully understand their functionality and importance (Dellinger, 2020). For instance, pollination and use of long spurred flowers does not necessarily correspond to long-proboscid visitors (Vlašánková et al., 2017), or that hummingbird visits are driven by nectar reward rather than other floral traits (Maruyama, Oliveira, Ferreira, Dalsgaard, & Oliveira, 2013). Furthermore, the observed trait configuration does not need to be an adaptation to pollinators alone. Floral antagonists exert negative selection pressures on floral traits, like flower size or the number of flowers on a plant, counteracting pollinator mediated selection pressures (Gélvez-Zúñiga, Teixido, Neves, & Fernandes, 2018). For example, it is still questionable to which extent red flowers are an adaptation to bird vision, or a defence against nectar thieving bees (Chittka & Waser, 1997; Rodríguez-Gironés & Santamaría, 2004; Wester et al., 2020) and if this role shifts spatially (Z. Chen, Niu, Liu, & Sun, 2020). Moreover, from a methodological point of view, if we acknowledge that individual traits largely differ in their effects on plant-pollinator interactions (e.g. Maruyama et al., 2013; Schmid et al., 2015), and that their synergic effects are important (Fenster et al., 2015), then the original category-based ordinations and classifications of individual traits do not seem to be the best expression of the real situation in nature (Abrahamczyk et al., 2017; Fig. 1). Hence, new methods are needed to reflect more complex interactions among predictors (i.e. floral traits) and to assess their importance for pollinators (Cutler et al., 2007; Dellinger, Chartier, et al., 2019; K. A. Johnson, 2013; Pichler, Boreux, Klein, Schleuning, & Hartig, 2020).

**Fig. 1.**
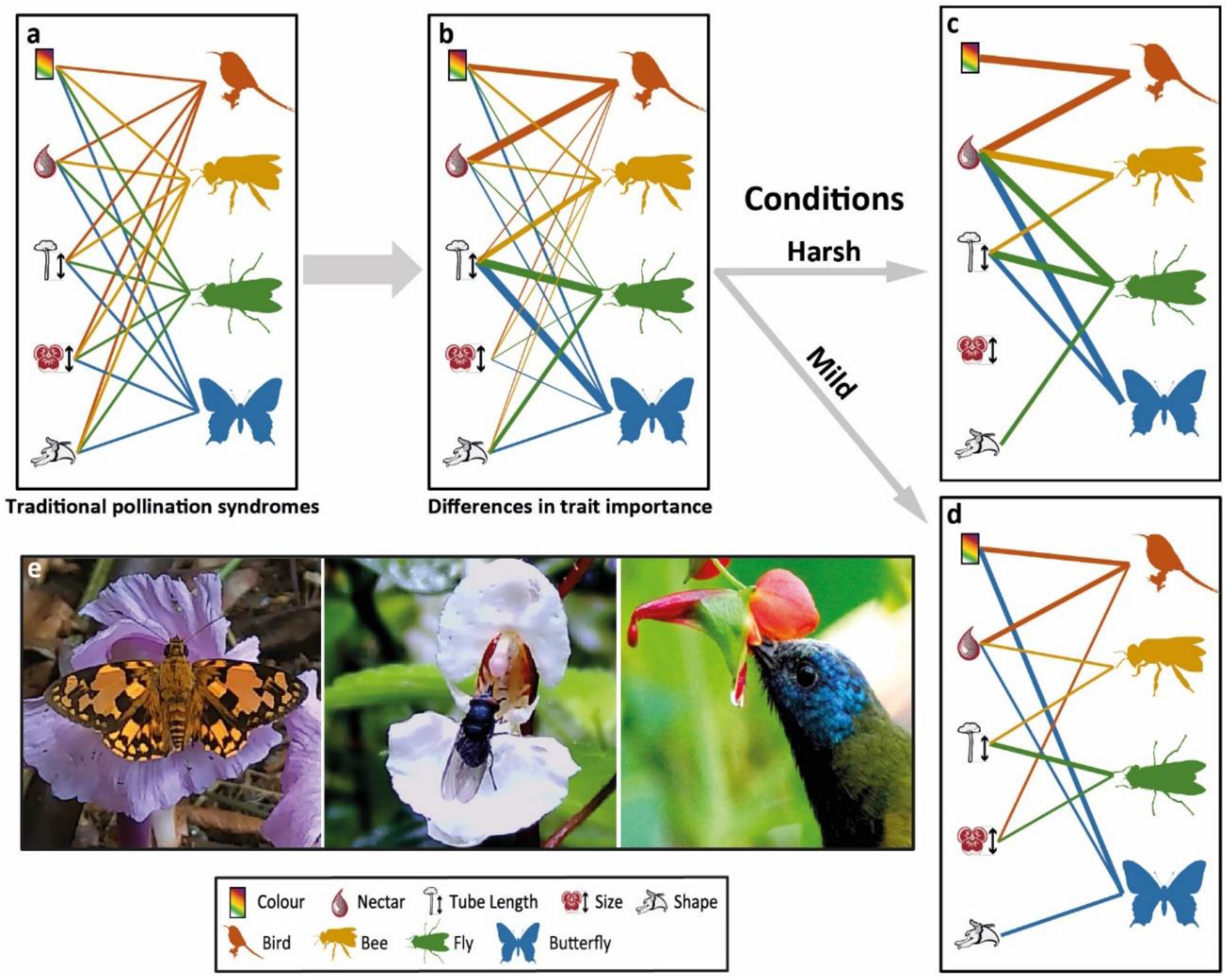
Conceptual figure. a) showing the traditional pollination syndrome view on the affiliation of plant traits and individual pollinator groups. b) floral traits differ in their relative importance for individual pollinator groups. c and d) The relative importance of individual floral traits differs between harsh and mild conditions: along the elevational gradient and between seasons. e) Examples of flower visitors found on Mount Cameroon (First two pictures are screenshots from the video recordings, the last picture was made by Š. Janeček). Note that relationships shown here do not reflect the real situation and are purely meant to conceptualise the hypotheses.

The second problem is that the role of individual floral traits and the related selection pressures can vary in space and time (Abrahamczyk, Kluge, Gareca, Reichle, & Kessler, 2011; Albrecht et al., 2018; González et al., 2009; Hawkins & Devries, 2009; Mayr et al., 2020; J. Mertens et al., 2020; Fig. 1c-d), which can make the formation of general conclusions regarding the ability of traits to predict pollinators even more challenging. However, understanding these spatiotemporal patterns can be crucial for revealing the role of floral traits in shaping plant-pollinator interactions.

Elevational gradients in seasonal ecosystems feature high adaptive trait differentiation and are thus an ideal place to study possible spatiotemporal variability in the relative importance of individual floral traits. Elevational gradients allow us to observe substantial changes in abiotic and biotic conditions (Girardin et al., 2014; McCain & Grytnes, 2010), including changes in taxonomical and functional diversities of plants and pollinators (Albrecht et al., 2018; Cuartas-Hernández, Moreno-Betancur, Gibernau, Herrera-Palma, & Hoyos-Serna, 2019; Janeček, Bartoš, & Njabo, 2015), as well as their interactions (Olesen & Jordano, 2002; Ramos-Jiliberto et al., 2010). Shifts in occurrence and/or abundances of individual floral traits have also been reported, their role in shaping biodiversity, however, remains unclear (Sun, Gross, & Schiestl, 2013). In addition, these elevational patterns can also differ seasonally (Maicher et al., 2020).

The importance of individual floral traits can be related to pollinator requirements and pollinator-community organisation. At higher elevations, pollinators have greater energetic requirements due to lower temperatures (Classen et al., 2015) or lower air pressure which hinders flight (Feinsinger, Colwell, Terborgh, & Chaplin, 1979). In turn, this can increase the importance of floral traits related to rewards (e.g. nectar production). Nectar production can increase in importance under less suitable seasonal conditions, such as during the wet season in humid tropical forests (Cruden, 1972; Janeček et al., 2015). This might be connected with the increasing prevalence of larger pollinators during wet seasons, such as nectarivorous birds, whose flight is less effected by rainy conditions compared to insects (Cruden, 1972). In contrast, Robinson & Wilson (1998) argued that according to the optimal foraging theory, increased resource availability could lead to greater niche partitioning, and thus potentially greater pollinator predictability by floral traits. We found increased resource availability at lower elevations (unpublished results), thus traits related to floral advertisement (e.g. colour or scent) could be of greater importance there. In seasons with less resources we can expect higher specialisation (Souza et al., 2018) and consequently, an increased ability to predict pollinators using floral traits.

To reveal the spatiotemporal variability in predictive power and relative importance of individual floral traits, we collected data on plant-pollinator interactions and floral traits on the community level, at four rainforest elevations along an elevational gradient on the highest West African mountain, Mount Cameroon. Our data was sampled in distinct wet and dry seasons, with extreme rains in the wet season (<2,000 mm monthly) and almost no rain during the dry season (Maicher et al., 2020). We test the following hypotheses: 1) The relative importance of individual floral traits differs among individual pollinator groups (Fig 1b). 2) There are environmentally driven changes in trait distribution and importance: 2.1) There is differentiation within floral trait distribution along the elevational gradient and between seasons. 2.2) Changes in the ability of traits to predict pollinators are found in harsh versus mild environmental conditions, i.e. along the elevational gradient and between seasons (Fig1c-d).

## Results

Based on 24h video recordings of flowers of 121 plant species with a complete set of floral traits, we identified 13,024 individual interactions of our selected potential pollinator groups, based on their contact with reproductive organs (See Material and Methods, and Supplementary Table 1). Using an extensive floral trait database (Supplementary Table 2) for all co-occurring flowering species at our focal sites, we were able to identify the importance of each trait in predicting pollinator groups across elevation and seasons. Using Random Forest models (RF), we identified the six most important floral traits for shaping plant-pollinator interactions on Mount Cameroon based on their Gini indices: Sugar per flower, Size, Shape, Tube Length, Colour and Flower position (Table 1). Nevertheless, we found seasonal and elevational shifts in trait importance for specific functional groups of primary pollinators when using individual RF analyses per season and elevation (Table 2).

**Table 1.** Ten floral traits used in the Random Forest analyses ranked by their importance (mean decrease in Gini index and mean decrease in accuracy) in distinguishing the eleven potential primary pollinator groups averaged for the 100 RFs with 500 trees each. Note that some pollinator groups were not found to be primary pollinators of any plant species in any season or elevation and were therefore excluded.

|  | Mean<br>Decrease<br>Accuracy | Mean<br>Decrease<br>Gini | Beetle | Hoverfly | Other Fly | Carpenter<br>bee | Honeybee | Wasp | Other Bee | Butterfly | Other Moth | Bird |
| --- | --- | --- | --- | --- | --- | --- | --- | --- | --- | --- | --- | --- |
| Sugar per Flower | 5.43 | 20.37 | 2.33 | 4.59 | 7.34 | 3.6 | 1.9 | -1.76 | -0.2 | -0.2 | -6.68 | 7.46 |
| Size | 5.17 | 19.29 | 3.15 | 0.28 | 4.24 | 1.34 | 1.1 | -0.65 | 3.5 | 4.88 | 1.12 | -2.74 |
| Shape | 9.08 | 14.66 | 3.24 | 2.6 | 1.12 | 0.33 | 3.82 | -0.18 | 3.51 | 2.79 | 6.92 | 2.13 |
| Tube Length | 6.39 | 12.64 | 5.13 | 1.93 | 2.22 | 1.9 | -0.49 | 2.41 | 5.54 | 0.49 | 4.65 | -3.62 |
| Colour | 9.36 | 11.92 | -2.9 | 1.67 | 8.76 | 0.63 | -0.76 | 0 | 8.36 | -0.6 | 3.55 | -1.45 |
| Flower Position | 0.36 | 7.66 | -0.2 | 3.12 | 0.26 | -1.89 | -4.51 | -1.5 | 0.12 | -4.6 | 4.33 | 0.96 |
| Odour Strength | 6.01 | 3.63 | 0.19 | -0.9 | -0.59 | 2.34 | 2.82 | 1.42 | 6.33 | -3.24 | 5.31 | -3.19 |
| Anther Position | 2.33 | 3.34 | 1.71 | 2.7 | 3.17 | 1 | -0.33 | 1 | 0.63 | -1.31 | 2.26 | -3.32 |
| Nectar Guides | 0.16 | 2.61 | 4.62 | 0.64 | -1.35 | -1.42 | -0.52 | 1.42 | 0.25 | -0.9 | -1.42 | -3.17 |
| Symmetry | 2.03 | 2.18 | 0.69 | -0.36 | 0.45 | -1.67 | -0.69 | -1.9 | 3.48 | -2.07 | 2.87 | -2.13 |

**Table 2.**
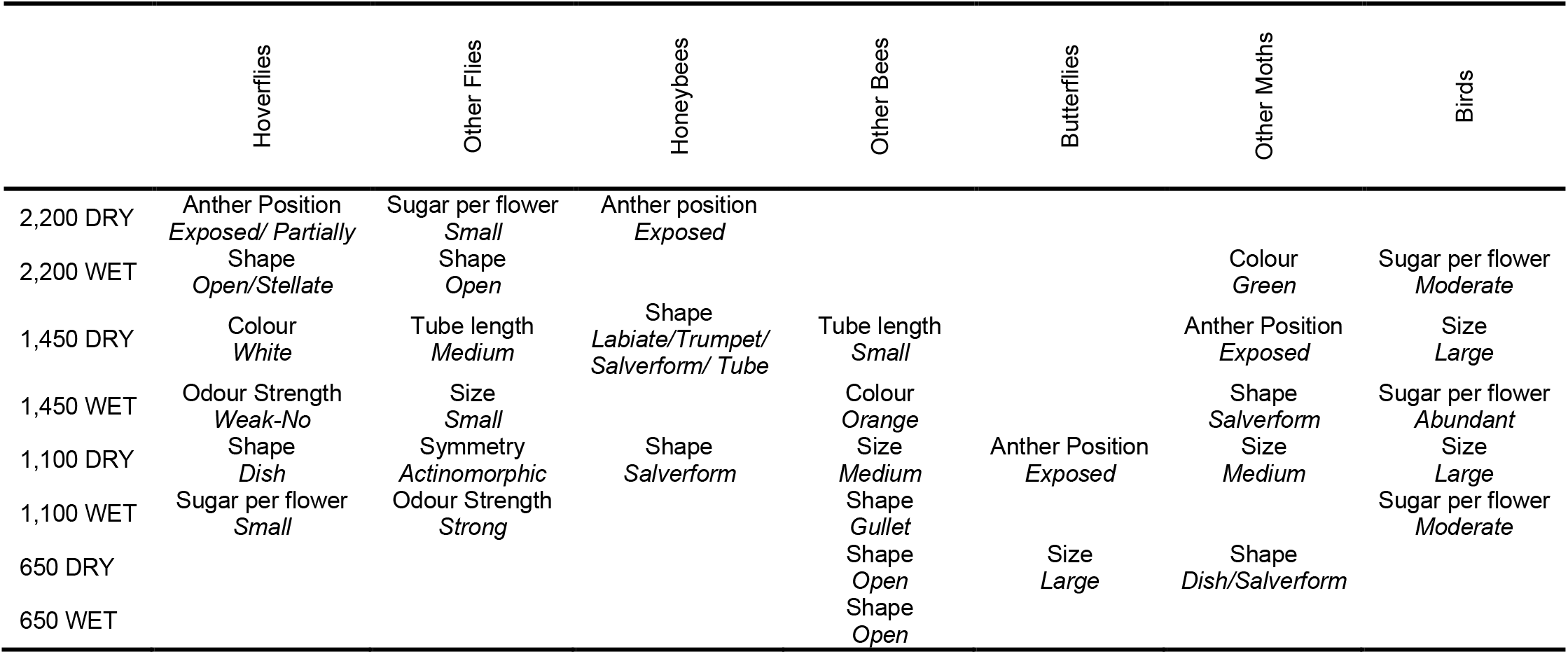
Most important trait per primary pollinator group according to the Random Forest analyses per season and elevation. The specific trait characteristics are given below each trat and are based on the highest occurrence of said traits in our dataset, for each specific season and elevation. For the continuous traits (size, tube length, or sugar per flower), the table shows whether the trait value is small, medium/moderate, or large/abundant compared to the rest of the database. Three groups were excluded from the table since they only were significant in a single season at a single elevation: Beetles (1,100m Dry: Shape *Open*), Carpenter bees (1,100m dry: Odour strength *Strong*) and Wasps (1,100m wet: Tube length *Small*).

These traits were also shown to be of similar importance when considering secondary pollinators (Supplementary Table 3). These machine learning algorithms can also be used for predicting pollinators based on floral trait combinations. For both primary and secondary pollinators we found a high prediction success when comparing individual pollinator groups (Table 3) and, along season and elevation (Table 4). Nevertheless, secondary pollinators (i.e. the second most frequent visitor touching reproductive organs) in general were less well predicted (Table 3 and 4).

**Table 3.**
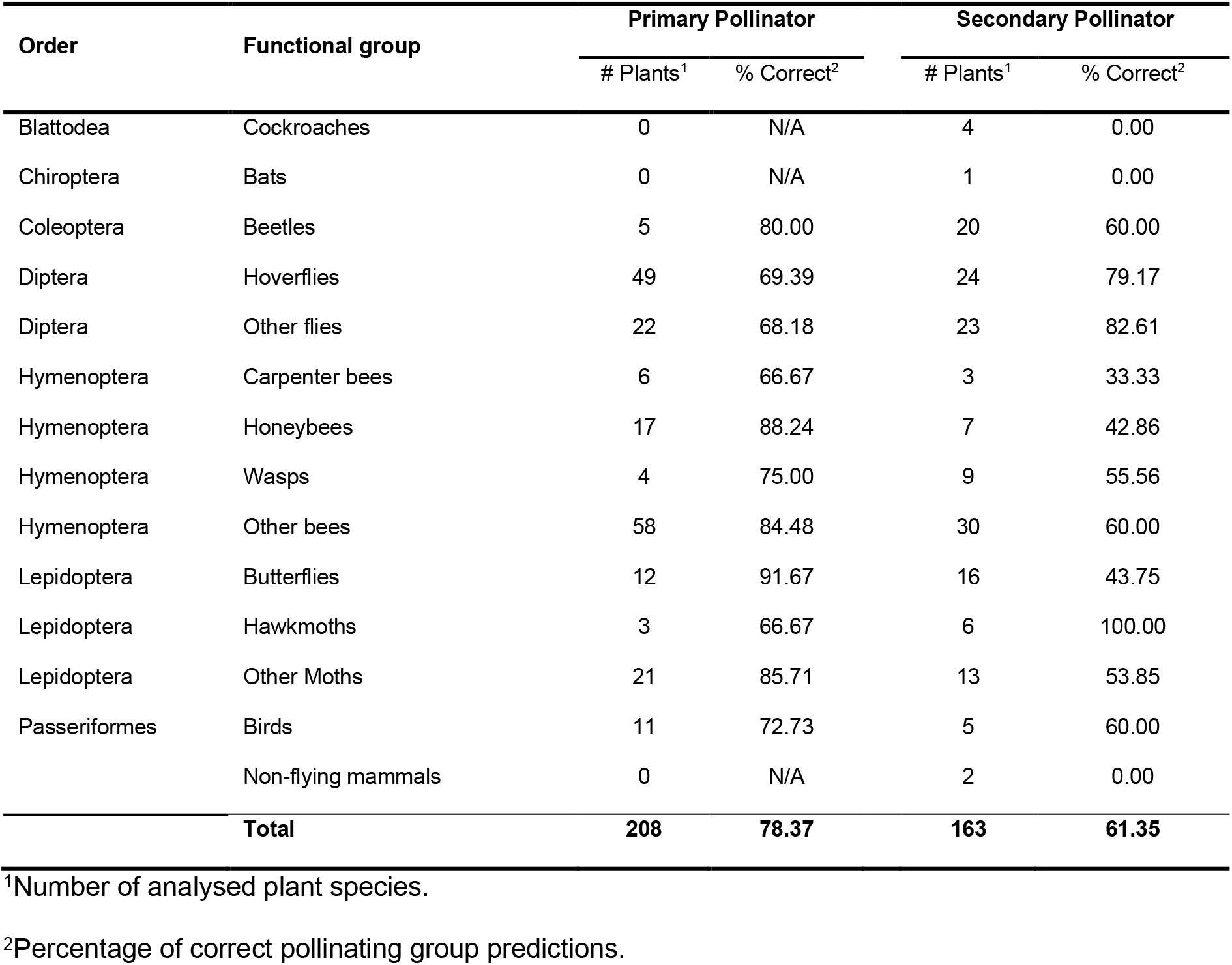
Predictions of potential pollinators per potential pollinator group by comparing the trained Random Forest model based on the floral traits with the actual primary or secondary pollinator found visiting flowers of particular plant species. N/A means the pollinator group did not occur as primary or secondary pollinator of any plant species in the given elevation and season.

**Table 4.** Predictions of potential pollinator groups per season and elevation by comparing the trained Random Forest model with the actual primary or secondary pollinator found visiting flowers of particular plant species.

| Elevation | Season | Primary Pollinator |  | Secondary Pollinator |  |
| --- | --- | --- | --- | --- | --- |
|  |  | # Plants<sup>1</sup> | % Correct<sup>2</sup> | # Plants<sup>1</sup> | % Correct<sup>2</sup> |
| 2,200m | DRY | 18 | 100.00 | 15 | 86.67 |
|  | WET | 19 | 84.21 | 12 | 75.00 |
| 1,450m | DRY | 24 | 70.83 | 21 | 42.86 |
|  | WET | 26 | 80.77 | 21 | 71.43 |
| 1,100m | DRY | 39 | 69.23 | 32 | 59.38 |
|  | WET | 24 | 70.83 | 21 | 57.14 |
| 650m | DRY | 37 | 86.49 | 26 | 53.85 |
|  | WET | 21 | 71.43 | 15 | 60.00 |
| **Total** |  | **208** | **78.37** | **163** | **61.35** |

<sup>1</sup>Number of analysed plant species.
<sup>2</sup>Percentage of correct pollinating group predictions.

Following the identification of the most important traits we used canonical correspondence analysis (CCA) to unveil any potential patterns in the trait distribution between dry and wet season and among particular elevations (Fig. 2a;F=1.2, P=0.044). The CCA shows that closed (e.g. gullet) and pendant/horizontal flowers are more common in the wet season, whilst the distribution of sugar per flower seems closely related to higher elevations (Fig. 2a). To test trait importance at a finer scale; per season and elevation, we performed eight separate forward selection CCA’s, with the five most important trait values (Fig. 2b; for significance see Supplementary Table 4). The models revealed an increase in the explained variability towards higher elevations and from dry to wet season, with the exception of the wet season at 2,200m, thus showing the importance of floral traits in distinguishing potential pollinators in harsher conditions. Additionally, we found that during the dry season, tube length and floral size were of greater importance for attracting pollinators, while in the wet season, red and orange flowers were most attractive (Fig. 2b). We also detected changes in trait importance with elevation, but these patterns were less apparent (Fig. 2b). Additionally, the variation explained by individual floral traits was greatest towards the highest elevations, and in the wet season generally (except for the highest elevation). Nevertheless, it must be noted that no statistically significant traits were found in the wet season at the two lowest elevations.

**Fig. 2.**
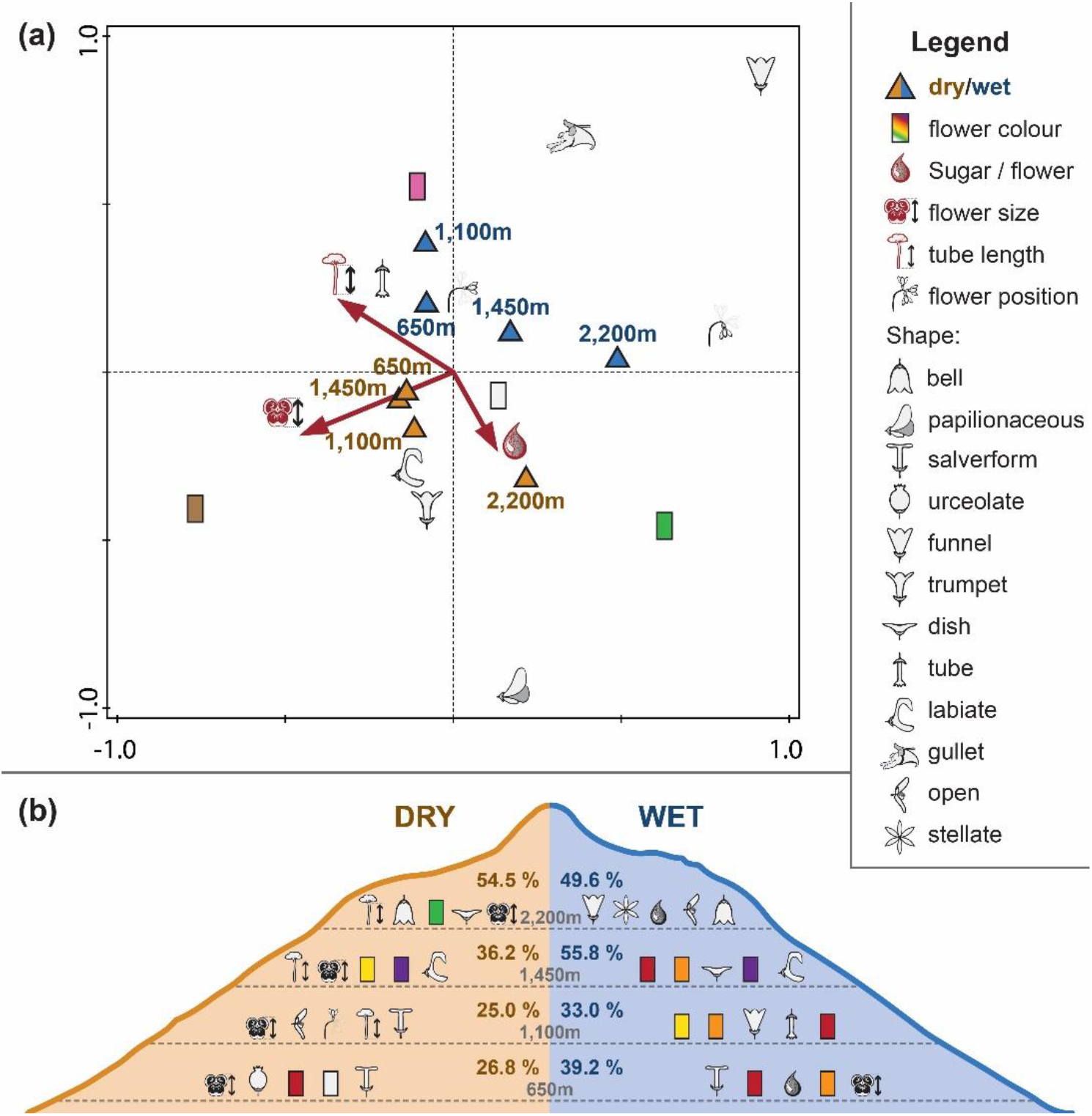
a) CCA ordination diagram visualising the distribution of individual floral traits between season and along elevation (listed here as m a.s.l.). b) Summarised outcomes of individual CCA’s of floral trait importance per elevation and season. Visualising the five floral trait characteristics with the highest ability to predict primary pollinators. The traits are sorted from left to right according to decreasing importance. Percentages in the middle states the variation explained by each model. The flower position symbol depicts one of the three possible flower orientations (Upright, Horizontal or Pendant).

## Discussion

We demonstrated that the relative importance of floral traits differs among the individual pollinator groups, and that there is spatiotemporal variability in floral trait distribution and importance (i.e. ability to predict pollinators) in the rainforest of Mount Cameroon.

The role of specific floral traits for flower visitors has been widely studied in numerous pollinator groups (e.g. Junker et al., 2013; Stang, Klinkhamer, Waser, Stang, & Van Der Meijden, 2009), with for example flower form and symmetry being important traits for flies (Lázaro, Hegland, & Totland, 2008). Compared to bees, flies are more common pollinators in the harsher conditions of higher elevations (Willmer, 2011). Traits related to this pollinator group (e.g. open/dish flowers) were found to be consistently important at the highest studied elevation, especially in the wet season (Fig. 2, Table 2). For bees, floral shape had the most predictive power. Open flowers were preferred in the lowest elevation (independent of the season), which might be explained by the flower offering a landing platform for bees, or by having exposed anthers, thus allowing for easy pollen collection. The huge variety in bee species, morphology and reward preferences (e.g. pollen, nectar and resin) allows them to exploit a wide range of floral designs (Ollerton, 2017; Willmer, 2011). This can also be seen in our study where flower shapes with a floral tube (salverform, tubular, trumpet) were the most important for honeybees at middle elevations (1,100m and 1,450m; Table 2). In a global analysis of pollination syndromes (Ollerton et al., 2009), bee and fly pollinated plants were predicted more accurately compared to other syndromes, while other studies did not show such patterns (K. A. Johnson, 2013). However, focusing only on the single most frequent visitor may not accurately depict true pollination, since visitation might be clouded by generalist visitors (Dellinger, 2020; Junker & Blüthgen, 2010; Ollerton et al., 2009; Padyšáková, Bartoš, Tropek, & Janeček, 2013; Rosas-Guerrero et al., 2014). Additionally, plants with morphologically generalised flowers are prone to this mismatch (Bartoš et al., 2015).

We found the occurrence of closed flowers plays a bigger role at higher elevations, unlike Jacquemyn, Micheneau, Roberts, & Pailler (2005), who showed a sharp decrease of species displaying floral traits related to hawkmoth (long spur) or fly pollination (small or no spur) in orchid species along the elevational range of Réunion Island. We found the opposite pattern for floral traits considered important for fly pollination, since open/dish and dull-looking (green/white colours) flowers were among the most important traits found at higher elevations (Fig. 2a-b, Table 2). These dull floral colours are often associated with hawkmoths (Faegri & van der Pijl, 1979; Willmer, 2011). However, tube length and scent have been shown to be more important traits for attracting hawkmoths (Willmer, 2011). We found shifts in the importance of corolla tube length only between dry and wet season and not along elevation (Fig. 2b).

Seasonality also had a strong effect on floral trait occurrence and their ability to predict individual pollinator groups. The higher occurrence of closed flowers (gullet, funnel and tube) and downwards-facing flowers in the wet season might be explained by their role as shelter for visitors, limiting the dilution of nectar (Aizen, 2003; Dafni, 1996; Pacini & Nepi, 2007) or washing and/or damaging of pollen (Huang, Takahashi, & Dafni, 2002; Pacini & Franchi, 1984). Additionally, rainfall affects the flight ability of potential pollinators through increased thermoregulatory costs, rain avoidance or as environmental noise (Lawson & Rands, 2019). For birds and bats it has been shown that rainfall increases the energy cost of flight (Aizen, 2003; Ortega-Jimenez & Dudley, 2012; Voigt, Schneeberger, Voigt-Heucke, & Lewanzik, 2011). This effect is even greater for smaller insect visitors whose activity can be partially reduced or even completely impeded, by directly damaging them and reducing their abundance (C. Chen, Harvey, Biere, & Gols, 2019; Kishimoto-Yamada & Itioka, 2015; Maicher et al., 2020, 2018; Struck, 1994). Similarly, the higher energetic requirements in upper elevations resulting from lower temperatures, is consistent with the increased nectar production per flower with elevation. Nectar production was a consistently good predictor of pollinators in both the highest (2,200 m) and lowest (650m) elevations during the wet season. In agreement with Aizen (2003) and Voigt *et al.* (2011), nectar sugar production was found to be especially important during the wet season for the most energy demanding pollinator group, birds. Since this group is more capable of dealing with harsher conditions (Cruden, 1972), floral traits associated with birds are expected to stand out. According to the individual RF analyses (Table 2) nectar sugar per flower and size were more important for bird pollination. However, red flower colour, which is generally strongly associated with bird pollination (Chittka & Waser, 1997; Rodríguez-Gironés & Santamaría, 2004; Wester et al., 2020), was found to be the third most important explanatory trait for birds in our study (Table 1) and particularly important in the wet season (Fig 2b).

Furthermore, we also show spatiotemporal differences in the general ability of traits to predict pollinators. We found an increase in explained variation towards the higher elevations, with the notable exception of the wet season at 2200m. For now, we can only speculate that this might be due to higher pollen limitation in upper elevations (and thus stronger selection pressures; Ashman et al., 2004; Knight et al., 2005) or that the plant and pollinator interactions, abundance and richness effects can be of influence.

Floral traits were better predictors of pollinators in the wet season, with the exception of the highest elevation. Souza *et al.* (2018) suggested that differences in resource availability in individual seasons can result in differences in interspecific competition among pollinators and niche overlaps. This suggestion was based on their observations from grassland and shrubby vegetation in the Pantanal and Cerrado ecosystems in Central Brazil, where the dry season was associated with a lack of resources and thus potentially higher levels of competition. In contrast, we found nectar production per hectare on Mount Cameroon was several times lower in the wet season (unpublished results). Nevertheless, we assume that, similar to Souza *et al.* (2018), higher competition and reduced niche overlap in the period of resource shortage can play a role, and these processes can result in higher plant specialisation and therefore greater predictability of pollinators by floral traits.

As we have shown, not all floral traits are equally important for individual pollinator groups and even traits that were not associated with traditional syndromes were found to be potentially important for predicting pollinators (Dellinger, 2020). From a methodological standpoint, machine learning approaches (such as RF) offer an avenue for dealing with such shifts in individual trait importance, increasing our predictive power. As a result, we have begun to see its application in several studies on trait selection, matching and syndrome testing (Dellinger, Chartier, et al., 2019; K. A. Johnson, 2013; Pichler et al., 2020). However, the necessity to remove incomplete and collinear traits when applying this trait based method has also been apparent (Dellinger, Chartier, et al., 2019; K. A. Johnson, 2013; Pichler et al., 2020). Regardless, even without a priori floral trait selection, the robustness of RF seems to allow for an ecologically realistic inference of pollinator predictability (Pichler et al., 2020).

## Conclusion

Our results showed the importance of floral traits in shaping plant-pollinator interactions in the understudied Afrotropics, with traits being more important towards the harsher conditions of higher elevations and wet seasons. These shifts in floral trait dependence between season and elevation show the importance of including spatiotemporal factors within pollination studies. The pollination syndromes will, in turn, vary in their ability to predict primary pollinators for different conditions and for the different pollinating groups. Additionally, shifts in the importance of specific discriminative traits for individual pollinator groups, together with spatiotemporal differences within these groups, suggests that following a complex predetermined list of equally important traits can be problematic for classifying potential pollinators. Using Random Forest models to categorise plants by floral traits and predict potential interactions enabled us to abandon the traditional pollination syndrome approach. Although the concept of pollination syndromes is still highly interesting, we propose stepping back and first unpacking the pollination syndromes into individual traits and improving our trait-based understanding of plant-pollinator interactions at the community level under different spatiotemporal and environmental conditions (Schmid et al., 2015).

## Material and Methods

### Study locality

This study was carried out on Mount Cameroon, Southwest Region of Cameroon (4.203°N and 9.170°E), the highest mountain in western and central sub-Saharan Africa (4,095 m a.s.l; Cable & Cheek, 1998). It represents an important biodiversity and endemism hotspot due to its location in the Cameroon Volcanic line (Gulf of Guinea Highlands) on the border of the Congo and Guinean bioregions, offering a wide range of habitats (Cable & Cheek, 1998; Maicher et al., 2020, 2018; Sosef et al., 2017).

We focused on the continuous elevational gradient of pristine rainforests, from lowland (±650 m a.s.l.), sub-montane (±1100 and ± 1450 m a.s.l.) to montane (± 2200 m a.s.l.) forests at the natural timberline on the southwestern slope of the mountain (For more details see Maicher et al., 2020). The region is known for its strong seasonality with annual precipitation exceeding 12,000 mm at the lower elevations in proximity to the Atlantic ocean with monthly precipitation in the wet season (June to September) of over 2,000 mm and almost no rainfall during the dry season (Maicher et al., 2020, 2018). At each studied elevation, six transects of 200m long and 10m wide (5m on both sides of the transect-line) were established at least 100m apart to cover the heterogeneity of the forest. Data sampling was performed along these transects; however, the search area also included the surrounding vegetation when there were insufficient replicates for a particular plant species.

### Floral trait measurements

Following Ollerton and Watts (2000) and Ollerton *et al.* (2009) we selected floral traits important for potential pollinators (Supplementary Table 2). Morphometric traits were measured by an electric calliper, visual and olfactory traits were recorded by the observer. Up to five individuals of all 121 plant species were measured.

Quantification of nectar sugar production was done by covering flowers with mesh bags for a 24-hour period. Nectar from individual flowers with high nectar production was extracted following Bartoš *et al.* (2012) using capillary tubes. The nectar concentration was measured using a Pal-1 (Atago co.) pocket refractometer to calculate the amount of sugar per flower. For low nectar-producing flowers, we washed the flowers using a Hamilton syringe with filtered water, added ethanol to the samples and boiled it for 15 minutes to deactivate enzymes. Nectar samples were dried in the laboratory, where they were transferred into constant volumes. The concentrations of individual sugars were measured with High-Performance Liquid Chromatography (HPLC) using the ICS-3000 system (Dionex) with an electrochemical detector and CarboPac PA 1 column. Our main dataset includes more than 121 plant species, but we excluded species that lacked complete floral trait measurements (morphometric or nectar) from our analyses.

### Visitor recording

The flowering plant species along the set transects were recorded for a 24-hour period using security cameras (VIVOTEK IB8367T with IR night vision; for more information on the methodology see Klomberg et al., 2019; Mertens et al., 2020, 2018) to detect and identify flower visitors to 13 functional groups of pollinators. We recorded flowers at all vegetation strata from understory to canopies (reached using tree climbing methods). The functional groups were defined following the common pollination syndrome groups (Birds, Bats, Flies, Bees, Wasps, Butterflies, Hawkmoths, Other moths, Non-flying-mammals (Willmer, 2011). Bees were split into Honeybees, Carpenter bees (as the large representative of the recently defined “buzz pollination syndrome; De Luca & Vallejo-Marín, 2013) and Other bees to better reflect floral reward usage. Flies were split into Hoverflies and Other flies. Cockroaches were also considered as potential pollinators (Mertens et al., 2018; Vlasáková, Pinc, Jůna, & Kotyková Varadínová, 2019). Only visitors seen touching plant reproductive organs (anthers or stigmas) were considered as potential pollinators (called pollinators elsewhere in our paper) and included in our analyses (Padyšáková et al., 2013). The concept of most effective pollinator has been treated elsewhere as the product of visitation frequency or pollen transfer efficiency (e.g. Ashworth et al., 2015; Fenster et al., 2015; Rosas-Guerrero et al., 2014). However, due to the scale of this work we were not able to classify pollen transfer efficiency and therefore relied on contacts with reproductive organs as a proxy of pollination (similar to e.g. Biella et al., 2019).

### Statistical analyses

Following Dellinger, Chartier, et al. (2019) we used Random Forest models (RF; Breiman, 2001) to identify the most important floral traits differentiating potential primary and secondary pollinators. Secondary pollinators were included to account for the possibility that the most frequent visitor was not equivalent to the most effective pollinator (Ashworth et al., 2015; Barrios, Pena, Salas, & Koptur, 2016; Padyšáková et al., 2013). Additionally, RF’s were done for each season and elevation separately to explore differences in trait importance along season and elevation. We ran 100 permutations of RF, comprising from 200 to 500 trees each (decision made during model training) and two variables tested at each split (mtry). The importance of each variable (floral trait) for distinguishing the pollinator groups was ranked by the mean decrease in accuracy and the mean decrease in Gini index (the total decrease in node impurities from splitting on the variable) over all 100 RFs (Breiman, 2001; Dellinger, Chartier, et al., 2019). The most important trait per primary pollinator group for each elevation and season was extracted from these individual analyses. For each trait we listed the most common trait value found in our dataset for the separate pollinator groups. Additionally, to evaluate the ability of floral traits and trait combinations to predict potential pollinators, we used a trained full mountain RF model to predict the primary and secondary pollinators of plant species in each season and elevation. For the analyses we used the RANDOMFOREST package (Liaw & Wiener, 2002) in R 3.6.1 (R Core Team, 2019).

We used constrained ordination, namely canonical correspondence analysis (CCA), using the Canoco 5 software (ter Braak & Šmilauer, 2012), to analyse patterns in trait distribution and variance explained by the six most important traits found during the RF analyses, allowing us to further unravel differences in floral trait composition and their importance between seasons and with elevational changes. We included all visitors touching the reproductive organs in these analyses, i.e. not only primary and secondary pollinators. We did a CCA to detect patterns in the trait distribution between seasons and with elevation. Additionally, we performed eight separate forward selection CCA’s (4 elevations within both seasons) to reveal the 5 most important trait values for each elevation and season, together with the variation explained by them.

## Acknowledgments

We are grateful to Vincent Maicher, Luma Francis Ewome, Raissa Dywou Kouede, Esembe Jacques Chi, Karolina Hrubá, Hernani Oliveira, Zuzana Sejfová, Pavel Potocký, Pavel Kratochvíl and our Cameroonian field assistants/students for help in the field. All our video watchers, especially Ivan Šonský, Sailee Sakhalkar, Petra Janečková, Eliška Chmelová, and Marek Rybár, for their help in processing the video recordings. Furthermore, we would like to thank the staff of the Mount Cameroon National Park for their support. This study was performed with all the required authorisations of the Republic of Cameroon Ministries for Forestry and Wildlife and for Scientific Research and Innovation. Also, thanks go to Agnes Dellinger and Paolo Biella for answering statistical questions, and to Kryštof Chmel and Jordan Bishop for their feedback on a previous version of the manuscript, and to Conor Redmond for English corrections. Additionally, we would like to thank Jan Wieringa and other botanists for their help with plant identifications. Symbols in Figure 1 and 2 were adapted from Phylopic.org (pollinators) and theseedsite.co.uk (floral shapes).

Our research was funded by the Czech Science Foundation (16-11164Y and 18-10781S), Charles University (PRIMUS/17/SCI/8 and UNCE204069) and the Grant Agency of the Charles University (GAUK No. 356217).

