## Supplementary material 1-4 for "Spatiotemporal shifts in the role of floral traits in shaping tropical plant-pollinator interactions"

Supplementary Table 1 | Summary of the number of plant species recorded per season and elevation including total hours of recordings and number of visits per pollinator group. Note that in the Random Forest analyses only primary and secondary pollinators were used (i.e. first and second most frequent visitor touching plant reproductive organs), therefore some groups, although present here, might be missing from the RF analyses.

| Elevation | Season | Plants | Recording hours | Cockroaches | Bats | Beetles | Hoverflies | Other flies | Carpenter bees | Honeybees | Wasps | Other bees | Butterflies | Hawkmoths | Other moths | Birds | Non-flying mammals |
| --- | --- | --- | --- | --- | --- | --- | --- | --- | --- | --- | --- | --- | --- | --- | --- | --- | --- |
| 2,200m | DRY | 18 | 1910 |  |  | 37 | 219 | 136 |  | 1696 | 4 | 3 | 4 | 23 | 78 | 5 | 5 |
|  | WET | 19 | 1385 |  |  | 3 | 112 | 173 |  | 46 | 1 | 2 |  | 4 | 23 | 43 |  |
| 1,450m | DRY | 24 | 2139 | 6 | 39 | 68 | 183 | 130 | 31 | 489 | 6 | 141 | 161 | 27 | 206 | 18 | 3 |
|  | WET | 26 | 1745 |  |  | 5 | 836 | 196 |  | 5 |  | 218 | 25 | 3 | 122 | 98 | 1 |
| 1,100m | DRY | 39 | 2912 | 17 |  | 52 | 181 | 100 | 71 | 85 | 60 | 314 | 137 | 12 | 163 | 8 |  |
|  | WET | 24 | 1708 |  |  | 9 | 323 | 211 | 6 |  | 41 | 131 | 43 | 6 | 33 | 51 | 1 |
| 650m | DRY | 37 | 2651 | 2 |  | 29 | 19 | 12 | 9 | 117 | 33 | 1084 | 113 | 12 | 152 | 4 |  |
|  | WET | 21 | 1636 | 1 |  | 14 | 12 | 251 |  | 1 | 136 | 1001 | 65 | 10 | 27 |  |  |

Supplementary Table 2 | Floral traits measured in the field for analyses with explanations of the categories.

| **Floral trait** | **Categories** | **Note** | **Included in analyses** |
| --- | --- | --- | --- |
| Shape | Bell, Bowl, Dish, Funnel, Gullet, Labiate, Open, Papilionaceous, Salverform, Stellate, Trumpet, Tube, Urceolate | As defined by Faegri and van der Pijl (1979), Ramirez 2003 and Ollerton and Watts 2002) | Yes |
| Symmetry | Actinomorphic / zygomorphic |  | Yes |
| Size (cm) | Quantitative data | Zygomorphic flowers were measured along the horizontal and vertical axis | Yes |
| Flower position | Horizontal, Pendant, Upright, All | All is a separate group with flowers of a plant growing in all three directions | Yes |
| Anther position | Exposed, partially exposed, Hidden |  | Yes |
| Tube width (cm) | Quantitative continuous data | Measured at the middle of the tube. In some cases, it was measured at the opening of the tube | No, correlated to Tube length (Spearman’s correlation: 0.8) |
| Tube length (cm) | Quantitative continuous data |  | Yes |
| Odour strength | Strong, Weak-No | Measured by observer smelling, therefore categories were kept simple. | Yes |
| Colour | Blue, Brown, Green, Orange, Pink, Purple, Red, White, Yellow | Based on visual inspection of flower | Yes |
| Nectar guides | Presence/Absence | Based on visual inspection of flower | Yes |
| Sugar amount per flower (mg) | Quantitative continuous data | Adjusted amount of sugar per sample to amount per flower based on HPLC or direct measurement using a pocket refractometer | Yes |

Supplementary Table 3 | Ten floral traits used in the Random Forest analyses ranked by their importance (mean decrease in Gini index and mean decrease in accuracy) in distinguishing the eleven potential secondary pollinator groups averaged for the 100 RFs with 500 trees each. Note that some pollinator groups were not found to be secondary pollinators of any plant species and were therefore excluded from the analysis.

|  | Mean Decrease Accuracy | Mean Decrease Gini | Beetle | Hoverfly | Other Fly | Carpenter bee | Honeybee | Wasp | Other Bee | Butterfly | Hawkmoth | Other Moth | Bird |
| --- | --- | --- | --- | --- | --- | --- | --- | --- | --- | --- | --- | --- | --- |
| Sugar per Flower | -0.63 | 17.04 | -3.83 | 2.48 | 0.76 | 2.15 | -2.5 | -1.19 | 0.77 | -1.11 | -0.75 | -4.34 | -1.65 |
| Size | 1.76 | 16.84 | 1.91 | 3.52 | 0.39 | 3.8 | -1.71 | -2.33 | -1.24 | 0.83 | -0.07 | -1.49 | -1.17 |
| Shape | -0.68 | 13.03 | -0.9 | 0.53 | 0.69 | -1.34 | -1.88 | -1.11 | -1.36 | 2.13 | -0.59 | -1.83 | -0.99 |
| Tube Length | 0.62 | 12.75 | 1.13 | -4 | 2.52 | 2.46 | -1.9 | -2.43 | 0.02 | 1.39 | 5.67 | -0.43 | -1.61 |
| Colour | -2.93 | 9.85 | -0.84 | 1.08 | -3.45 | -1.9 | -1.27 | -0.85 | -2.59 | -1.96 | 3.92 | 0.09 | 0.07 |
| Flower Position | -1.55 | 6.36 | -3.53 | -2.09 | 1.69 | 2.66 | -1.34 | -0.29 | 1.11 | 3.32 | -1.4 | -3.08 | 0 |
| Odour Strength | 2.14 | 3.7 | 0.09 | 0.09 | 3.01 | -2.25 | -0.33 | -0.26 | -2.16 | 4.17 | 2.61 | -0.12 | -1.9 |
| Anther Position | 1.41 | 3.16 | 0.31 | 1.08 | 4.28 | 1 | -1 | -2.09 | -3.09 | 1.32 | -0.36 | -1.56 | -2.1 |
| Nectar Guides | 1.38 | 2.48 | -1.8 | 0.33 | 2.02 | 1 | 0 | 1 | 3.62 | -2.03 | -3.52 | -0.47 | 0 |
| Symmetry | 0.74 | 2.24 | 0.48 | -0.21 | -2.11 | 1.74 | 0 | -0.58 | 2.65 | 2.87 | -1.94 | 1.39 | -2.13 |

Supplementary Table 4 | Significance values of individual canonical correspondence

|  |  |  | **Shape** | | | | | | | | | **Flower position** | **Colour** | | | | | | **Size** | **Tube length** | **Sugar per flower** |
| --- | --- | --- | --- | --- | --- | --- | --- | --- | --- | --- | --- | --- | --- | --- | --- | --- | --- | --- | --- | --- | --- |
|  |  |  | *Bell* | *Salverform* | *Urceolate* | *Funnel* | *Dish* | *Tube* | *Labiate* | *Open* | *Stellate* | *Upright* | *Green* | *Orange* | *Purple* | *Red* | *White* | *Yellow* |  |  |  |
| **2,200 m** | **Dry** | Rank | 2 |  |  |  | 4 |  |  |  |  |  | 3 |  |  |  |  |  | 5 | 1 |  |
|  |  | Explained variation | 11 |  |  |  | 14.5 |  |  |  |  |  | 9.7 |  |  |  |  |  | 6.5 | 12.9 |  |
|  |  | F | 2.2 |  |  |  | 3.6 |  |  |  |  |  | 2 |  |  |  |  |  | 1.7 | 2.4 |  |
|  |  | P | 0.128 |  |  |  | **0.008** |  |  |  |  |  | 0.144 |  |  |  |  |  | 0.186 | 0.078 |  |
|  | **Wet** | Rank | 5 |  |  | 1 |  |  |  | 4 | 2 |  |  |  |  |  |  |  |  |  | 3 |
|  |  | Explained variation | 5.5 |  |  | 16.1 |  |  |  | 5.2 | 12.6 |  |  |  |  |  |  |  |  |  | 10.2 |
|  |  | F | 1.4 |  |  | 3.3 |  |  |  | 1.3 | 2.8 |  |  |  |  |  |  |  |  |  | 2.5 |
|  |  | P | 0.22 |  |  | 0.07 |  |  |  | 0.296 | **0.048** |  |  |  |  |  |  |  |  |  | 0.07 |
| **1,450 m** | **Dry** | Rank |  |  |  |  |  |  | 5 |  |  |  |  |  | 4 |  |  | 3 | 2 | 1 |  |
|  |  | Explained variation |  |  |  |  |  |  | 6.4 |  |  |  |  |  | 6 |  |  | 7.4 | 7.6 | 8.9 |  |
|  |  | F |  |  |  |  |  |  | 1.9 |  |  |  |  |  | 1.7 |  |  | 2 | 2 | 2.2 |  |
|  |  | P |  |  |  |  |  |  | **0.036** |  |  |  |  |  | **0.048** |  |  | **0.04** | **0.044** | **0.004** |  |
|  | **Wet** | Rank |  |  |  |  | 3 |  | 5 |  |  |  |  | 2 | 4 | 1 |  |  |  |  |  |
|  |  | Explained variation |  |  |  |  | 9.9 |  | 13.6 |  |  |  |  | 11.8 | 6.6 | 13.9 |  |  |  |  |  |
|  |  | F |  |  |  |  | 3.5 |  | 6.4 |  |  |  |  | 3.8 | 2.5 | 4 |  |  |  |  |  |
|  |  | P |  |  |  |  | 0.072 |  | **0.002** |  |  |  |  | **0.028** | 0.062 | **0.006** |  |  |  |  |  |
| **1,100 m** | **Dry** | Rank |  | 5 |  |  |  |  |  | 2 |  | 3 |  |  |  |  |  |  | 1 | 4 |  |
|  |  | Explained variation |  | 4.2 |  |  |  |  |  | 6.5 |  | 4.3 |  |  |  |  |  |  | 5.8 | 4.2 |  |
|  |  | F |  | 1.9 |  |  |  |  |  | 2.7 |  | 1.9 |  |  |  |  |  |  | 2.3 | 1.9 |  |
|  |  | P |  | **0.04** |  |  |  |  |  | **0.002** |  | **0.032** |  |  |  |  |  |  | **0.012** | **0.028** |  |
|  | **Wet** | Rank |  |  |  | 3 |  | 4 |  |  |  |  |  | 2 |  | 5 |  | 1 |  |  |  |
|  |  | Explained variation |  |  |  | 6 |  | 5.5 |  |  |  |  |  | 7.3 |  | 5.56 |  | 8.5 |  |  |  |
|  |  | F |  |  |  | 1.5 |  | 1.4 |  |  |  |  |  | 1.8 |  | 1.5 |  | 2.1 |  |  |  |
|  |  | P |  |  |  | 0.204 |  | 0.17 |  |  |  |  |  | 0.072 |  | 0.118 |  | 0.078 |  |  |  |
| **650 m** | **Dry** | Rank |  | 5 | 2 |  |  |  |  |  |  |  |  |  |  | 3 | 4 |  | 1 |  |  |
|  |  | Explained variation |  | 5.1 | 5.8 |  |  |  |  |  |  |  |  |  |  | 5.4 | 3.7 |  | 6.8 |  |  |
|  |  | F |  | 2.3 | 2.4 |  |  |  |  |  |  |  |  |  |  | 2.3 | 1.6 |  | 2.7 |  |  |
|  |  | P |  | **0.02** | 0.108 |  |  |  |  |  |  |  |  |  |  | **0.028** | 0.064 |  | **0.016** |  |  |
|  | **Wet** | Rank |  | 1 |  |  |  |  |  |  |  |  |  | 4 |  | 2 |  |  | 5 |  | 3 |
|  |  | Explained variation |  | 13.2 |  |  |  |  |  |  |  |  |  | 5.8 |  | 7.5 |  |  | 6.1 |  | 6.6 |
|  |  | F |  | 2.9 |  |  |  |  |  |  |  |  |  | 1.4 |  | 1.7 |  |  | 1.5 |  | 1.5 |
|  |  | P |  | 0.056 |  |  |  |  |  |  |  |  |  | 0.164 |  | 0.168 |  |  | 0.17 |  | 0.138 |

Analyses (CCA) per season and elevation. This table supplements Fig. 2B, it shows the significance of the ranked traits per season and elevation. Rank (1-5), explained variation (%), Pseudo-F and P value are displayed.
